# Factors associated with unsuppressed viremia in women living with HIV on lifelong ART in a multi-country cohort study: US-PEPFAR PROMOTE study

**DOI:** 10.1101/688945

**Authors:** Patience Atuhaire, Sherika Hanley, Nonhlanhla Yende-Zuma, Jim Aizire, Lynda Stranix-Chibanda, Bonus Makanani, Beteniko Milala, Haseena Cassim, Taha Taha, Mary Glenn Fowler

**Affiliations:** Makerere University-Johns Hopkins University (MU-JHU) Kampala, Uganda; Centre for the AIDS Programme of Research in South Africa (CAPRISA), Umlazi Clinical Research Site, Nelson R. Mandela School of Medicine, Durban, South Africa; Centre for the AIDS Programme of Research in South Africa (CAPRISA), Durban, South Africa; Johns Hopkins Bloomberg School of Public Health, Department of Epidemiology, Baltimore, MD, USA; University of Zimbabwe College of Health Sciences Department of Paediatrics and Child Health; Malawi College of Medicine-John’s Hopkins Research Project; UNC Project-Malawi; Perinatal HIV Research Unit (PHRU), Chris Hani Baragwanath Hospital, University of Witwatersrand, Johannesburg, South Africa; Johns Hopkins University, Departments of Pathology and Epidemiology, Baltimore, MD, USA

## Abstract

**Background:** Despite recent efforts to scale-up lifelong combination antiretroviral therapy (cART) in sub-Saharan Africa, high rates of unsuppressed viremia persist among cART users, and many countries in the region fall short of the UNAIDS 2020 target to have 90% virally suppressed. We sought to determine the factors associated with unsuppressed viremia (defined for the purpose of this study as >200 copies/ml) among African women on lifelong cART.

**Methods:** This analysis was based on baseline data of the PROMOTE longitudinal cohort study at 8 sites in Uganda, Malawi, Zimbabwe and South Africa. The study enrolled 1987 women living with HIV who initiated lifelong cART at least 1 year previously to assesses long-term safety and effectiveness of cART. Socio-demographic, clinical, and cART adherence data were collected. We used multivariable Poisson regression with robust variance to identify factors associated with unsuppressed viremia.

**Results:** At enrolment, 1947/1987 (98%) women reported taking cART. Of these, HIV-1 remained detectable in 293/1934 (15%), while 216/1934 (11.2%) were considered unsuppressed (>200 copies/ml). The following factors were associated with an increased risk of unsuppressed viremia: not having household electricity (adjusted prevalence rate ratio (aPRR) 1.74, 95% confidence interval (CI) 1.28-2.36, p<0.001); self-reported missed cART doses (aPRR 1.63, 95% CI 1.24-2.13, p<0.001); recent hospitalization (aPRR 2.48, 95% CI 1.28-4.80, p=0.007) and experiencing abnormal vaginal discharge in the last three months (aPRR 1.88; 95% CI 1.16-3.04, p=0.010). Longer time on cART (aPRR 0.75, 95% CI 0.64-0.88, p<0.001) and being older (aPRR 0.77, 95% CI 0.76-0.88, p<0.001) were associated with reduced risk of unsuppressed viremia.

**Conclusion:** Socioeconomic barriers such as poverty, not being married, young age, and self-reported missed doses remain key predictors of unsuppressed viremia. Targeted interventions are needed to improve cART adherence among women living with HIV with this risk factor profile.

## Introduction

Since 2012, the rapid scale up of the World Health Organization (WHO) option B+ strategy among pregnant or breastfeeding women living with Human Immunodeficiency Virus (HIV) has resulted in a substantial reduction in maternal morbidity and mortality, as well as incident pediatric HIV infections(1). Subsequently with the introduction of the Universal “Test and Treat” strategy, approximately 21.7 million people (including women) had access to combination Antiretroviral Therapy (cART) globally in 2017. Ensuring sustained adherence to and virologic suppression on cART is paramount in achieving the Joint United Nations Programme on HIV and AIDS (UNAIDS) 90-90-90 2020 Strategy in ending the epidemic by 2030 (2).

Barriers to achieving the UNAIDS 2020 strategy regarding the ‘third 90’ persist in sub-Saharan Africa in part due to suboptimal cART adherence(3). In the absence of viral resistance, HIV viral load assessment is the proxy for adherence, therefore the contributing factors to both adherence and viremia may overlap. The most common factors identified as being associated with decreased adherence include individual factors like younger age below 24 years, forgetting the dosing time, depression, and substance use. The predominant contextual issue remains stigmatization and disclosure [3–5]. Other factors such as length of time on ART, education, personal motivation to start ART, satisfaction with health worker information availed were conducive to adherence [6–8]. Additionally, studies have shown that women who initiate cART for their own health display better adherence as compared to women who initiate during pregnancy. Even then, adherence to cART in the post-partum period tends to wane. This may be attributed to less motivation to protect the child post-delivery and following cessation of breast feeding, as well as a possible break in transition from postnatal to general HIV care [4, 5].

Whereas the scale up of virologic monitoring in sub-Saharan Africa since 2013 has led to the availability of data regarding factors associated with virologic detectability, there is a paucity of literature as to which factors are most strongly assoicated with unsuppressed viremia among African pregnant women and mothers living with HIV. What is known to date is that virological detectabilty in resource-limited settings has been associated with the presence of comorbidities like Tuberculosis or psychiatric disease, higher pretreatment HIV RNA levels, repeat testers after suspected virologic failure and initiation of cART late in pregnancy(4–9). Thus the purpose of this analyses is to determine clinical and demographic risk factors associated with unsuppressed viremia among a well characterized cohort of women living with HIV originally in the PROMISE clinical trial at the time of their entry into the PROMOTE Study.

The PROMOTE study is an observational cohort study of mothers with HIV and their children who had participated in the IMPAACT 1077BF/1077FF PROMISE (Promoting Maternal and Infant Survival Everywhere) study at high enrolling sites in Zimbabwe, Malawi, Uganda and S. Africa. The PEPFAR-funded PROMOTE study presents a unique opportunity to assess longer term treatment outcomes among women randomized in the PROMISE trial to initiate varied antiretroviral (ARV) regimens during pregnancy for the purpose of preventing perinatal HIV transmission and subsequently transitioned to lifelong cART following disease progression or in response to the START study which showed clear benefit of universal ART in June 2015 (10). The PROMOTE study approach provides data from current public sector HIV care provision mixed with precise individualized clinical and laboratory data collected under trial settings. The vast majority of existing studies have assessed factors associated with viremia >1000 copies/ml, the WHO threshold for treatment failure. Emerging antriretroviral drug resistance has been known to occur from levels of 200 copies/ml or above, and use of this threshold eliminates most cases of apparent viremia caused by viral load blips or assay variability. We therefore sought to assess factors associated with viremia above 200 copies/ml in PROMOTE women at baseline.

## Materials and Methods

### Design

The PEPFAR-PROMOTE study is a five-year observational cohort of African women with HIV and their children previously enrolled in the PROMISE (Promoting Maternal and Infant Survival Everywhere) randomized trial(11). Commencing three months after the PROMISE trial closed-out in September 2016, 1987 mothers and their children were recruited from the high enrolling PROMISE sites. Enrollment to PROMOTE was completed in August 2017. The PROMOTE study is one of longest ongoing follow up epidemiologic multinational cohorts in sub-Saharan Africa.

### Setting and study populations

The PROMOTE study is being conducted at eight research sites in four African countries: MUJHU/Kampala (Uganda), Blantyre and Lilongwe (Malawi), Harare Family Care, Seke North and St. Mary’s (Zimbabwe), PHRU/Johannesburg and CAPRISA Umlazi/Durban (South Africa).

### Inclusion criteria

Women and children enrolled in the PROMISE trial from the 8 high enrolling African PROMISE sites described above, who were willing to provide informed consent to enroll and continue follow-up in the PROMOTE study.

### Exclusion criteria

Women who were unwilling to provide informed consent to continue follow-up in the PROMOTE study; or had plans to relocate permanently out of the catchment area during the cohort study period; or judged by the site team as having social or other reasons which would make it difficult for the mother/child pair to comply with study requirements.

### Enrolment study procedures for women

Mothers with HIV previously enrolled in the PROMISE study were re-enrolled after appropriate counseling and consenting in the PROMOTE study. At the enrolment visit, trained study workers administered sociodemographic and ART adherence questionnaires. A complete medical history and physical examination, including WHO clinical staging, was performed. Included in a holistic package of comprehensive counseling was the provision of study-specific antiretroviral adherence counseling. Enrollment laboratory evaluations included: viral load and CD4+ cell count. Viral load tests at the different sites are done using the COBAS TaqMan and Abbot assays with a lower limit of detectability at 20 copies/ml and 40 copies/ml respectively. All questionnaire responses and laboratory data were completed on designated case report forms (CRFs) by trained research site personnel. Samples were stored for future HIV drug resistance testing (blood) and cumulative drug levels testing (hair).

### Ethical considerations

The PROMOTE study was approved by all relevant institutional review boards (IRBs) in the U.S. and participating African research sites/countries. All women provided written informed consent to enroll and be followed up for the duration of the study with their children and agreed to provide study samples for protocol lab safety assays, as well as for storage of blood and hair samples.

### Statistical analysis

We analyzed baseline data from a multi-country cohort study to estimate the proportion of women living with HIV who had unsuppressed viremia (defined as viral load above 200 copies/ml) and to identify predictors of unsuppressed viremia. Fisher’s exact, Chi-square test of independence or Wilcoxon rank sum tests were used to test for an association between baseline characteristics and unsuppressed viremia. We used multivariable Poisson regression with robust variance to identify the predictors of unsuppressed viremia, and calculated prevalence risk ratios to measure the strength of an association between baseline characteristics and unsuppressed viremia. This was done in two ways (i) each predictor was fitted in the model, and (ii) multivariable model with all the predictors included in the model. Variables included in the multivariable analyses were chosen based on prior research on risk factors, biological plausibility, and previously identified clinical associations. Models (i) and (ii) were adjusted for the country variable to account for variation in geographical location of the research sites. Variables with increased missing data (>20% of observations missing per variable) and variables that were highly correlated were not included in the multivariable model. In an exploratory analyses, women with detectable viral loads were further stratified into the following thresholds (<50, 50-200, 201-1000 and >1000 copies per ml based on the varied thresholds by various HIV cART committees in resource rich and resource limited settings)(12, 13).

## Results

Overall, 1987 mothers were enrolled into the PROMOTE study, of whom 1947 (98%) women reported taking ART at the enrolment visit and HIV-1 viral load results were available for 1934. HIV-1 VL was above the limit of quantification in 293/1934 (15 %). A total of 216/1934 (11.2%) presented with an unsuppressed viremia above 200 copies/ml. Furthermore, among the 293 women with detectable viral load, 24 (8.2%) had VL below 50, 53 (18.1%) had VL between 50 and 200, 50 (17.1%) had VL between 201 and 1000, while 166 (56.7%) had VL above 1000 copies/ml as displayed in Fig 1.

**Fig 1.**
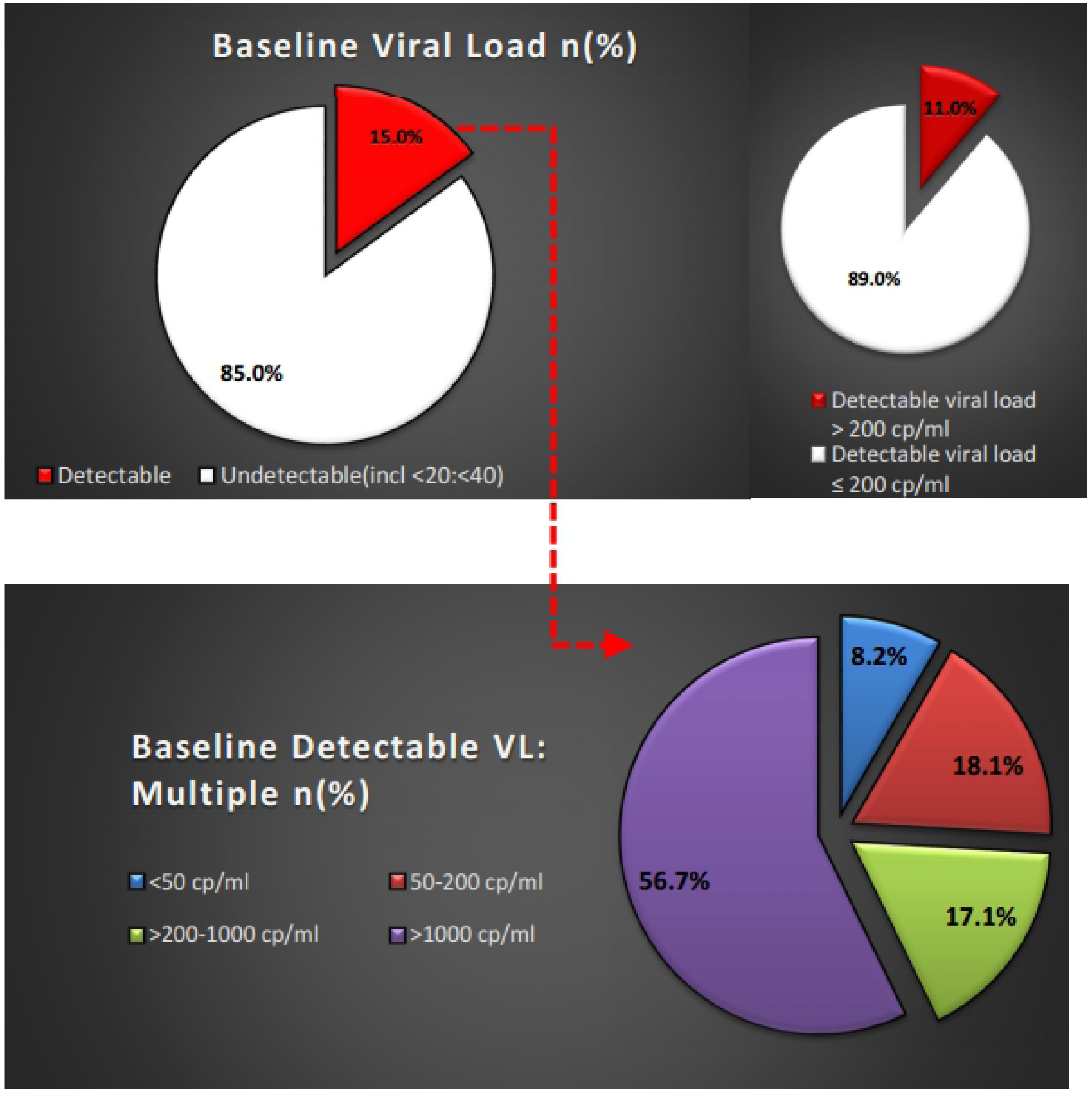
The proportions of viral suppression based on different thresholds of detectable viral load.

The individual and contextual baseline characteristics of the PROMOTE study have been reported elsewhere (11). Table 1 displays baseline characteristics stratified by viral load below and above 200 copies/ml. With the exception of employment status, HIV status disclosure, condom usage, all the baseline variables were associated with unsuppressed viremia >200 copies/ml (Table 1). Notably, the prevalence of unsuppressed viremia > 200 copies/ml was the highest (17.5%) in Malawi compared to other countries. Of note, 24% of women with recent hospitalization had unsuppressed viremia.

**Table 1:** Individual and contextual baseline characteristics.

| Variable | Viral load<br>≤200<br>copies/ml<br>(N=1718) | Viral load<br>>200<br>copies/ml<br>(N=216) | p-value |
| --- | --- | --- | --- |
| **Socioeconomic and demographic factors** |  |  |  |
| *Country, n (%)* |  |  | <.001 |
| Uganda | 319 (90.6%) | 33 (9.4%) |  |
| Malawi | 522 (82.5%) | 111 (17.5%) |  |
| Zimbabwe | 406 (90.6%) | 42 (9.4%) |  |
| South Africa | 471 (94.0%) | 30 (6.0%) |  |
| Age (years), median(IQR) | 31 (28 - 35) | 29 (25 - 33) | <.001 |
| Baseline CD4 cell count (cells/μL), median (IQR) | 852(674-1064) | 615(472-810) | <0.001 |
| *Marital status, n(%)* |  |  | 0.003 |
| Other (single, divorced, widowed, separated) | 325 (84.4%) | 60 (15.6%) |  |
| Married/regular partner | 1393 (89.9%) | 156 (10.1%) |  |
| *Employment, n(%)<sup>a</sup>* |  |  | 0.076 |
| Formal employment | 383 (90.5%) | 40 (9.5%) |  |
| Self-employment (small business) | 532 (86.5%) | 83 (13.5%) |  |
| Not employed/housewife | 802 (89.7%) | 92 (10.3%) |  |
| *Highest level of education, n(%)* |  |  | 0.011 |
| Secondary school completed or tertiary education | 1227 (90.0%) | 136 (10.0%) |  |
| Electricity in the premises, n(%) | 1209 (91.8%) | 108 (8.2%) | <.001 |
| *Tap water in the premises, n(%)* | 1139 (90.4%) | 121 (9.6%) | 0.004 |
| *Travel time from home to clinic, n(%)<sup>b</sup>* |  |  | 0.044 |
| Less than 30 minutes | 444 (92.3%) | 37 (7.7%) |  |
| 30-60 minutes | 757 (87.7%) | 106 (12.3%) |  |
| 1-2 hours | 389 (87.2%) | 57 (12.8%) |  |
| Greater than 2 hours | 127 (88.8%) | 16 (11.2%) |  |
| *Disclosed HIV status to partner, n(%)<sup>c</sup>* |  |  | 0.465 |
| Disclosed to partner | 1201 (90.2%) | 131 (9.8%) |  |
| No disclosure | 192 (88.5%) | 25 (11.5%) |  |
| *Partner's HIV status<sup>d</sup>* |  |  |  |
| Positive | 724 (90.2%) | 79 (9.8%) | 0.908 |
| Negative | 257 (89.9%) | 29 (10.1%) |  |
| *Condom usage during sex in last 3 months, n(%)<sup>e</sup>* |  |  | 0.811 |
| Always | 502 (89.8%) | 57 (10.2%) |  |
| Sometimes | 506 (88.6%) | 65 (11.4%) |  |
| Never | 265 (88.9%) | 33 (11.1%) |  |
| *Years on ART, median(IQR)* | 2 (1 - 2) | 1 (1 - 2) | <.001 |
| **Clinical factors** |  |  |  |
| Admitted to hospital in the past 3 months, n(%) |  |  |  |
| Yes | 22 (75.9%) | 7 (24.1%) | 0.036 |
| No | 1696 (89.0%) | 209 (11.0%) |  |
| Received TB treatment in the last 3 months, n(%) |  |  |  |
| Yes | 8 (88.9%) | 1 (11.1%) | 1.00 |
| No | 1710 (88.8%) | 215 (11.2%) |  |
| Presence of abnormal vaginal discharge in the last 3 months, n (%) |  |  |  |
| Yes | 80 (82.5%) | 17 (17.5%) | 0.047 |
| No | 1638 (89.2%) | 199 (10.8%) |  |
| Currently breastfeeding, n(%)<sup>b</sup> |  |  |  |
| Yes | 163 (84.9%) | 29 (15.1%) | 0.071 |
| No | 1554 (89.3%) | 187 (10.7%) |  |
| **ART related factors** |  |  |  |
| Never missed any doses since last visit, n(%)<sup>f</sup> | 1257 (91.8%) | 113 (8.2%) | <.001 |
| Number of days ARV doses missed in last four days, n(%)<sup>f</sup> |  |  |  |
| None | 1638 (90.6%) | 169 (9.4%) | <.001 |
| Awareness of dosing instructions, n(%)<sup>f</sup> |  |  |  |
| Yes | 826 (87.9%) | 114 (12.1%) | 0.017 |
| No | 842 (90.3%) | 90 (9.7%) |  |
| <sup>a</sup>2 missing data, <sup>b</sup>1 missing data, <sup>c</sup>amongst 1549 women with partners, <sup>d</sup> amongst women whom their partner's got tested and women knew their HIV status,<br><sup>e</sup>amongst women who report sexual activities, <sup>f</sup>amongst those who were on ART at enrollment |  |  |  |

The predictors of detectable viremia > 200cp/ml are shown in Table 2. Recent hospital admission and experiencing abnormal vaginal discharge in the last three months were associated with a 2.5-fold and almost 2-fold higher risk of detectable viremia respectively (adjusted prevalence risk ratio (aPRR) 2.48, 95% confidence interval (CI) 1.28-4.80, p=0.007; aPRR 1.88; 95% CI 1.16-3.04, p=0.010). In addition, the absence of socioeconomic factors such as electricity in the premises was associated with a 74% higher risk of detectable viremia (aPRR 1.74, 95% CI 1.28-2.36, p<0.001). Women who missed some of their ART doses were more likely to present with detectable viremia (aPRR 1.63, 95% CI 1.24-2.13, p<0.001). The most common reason for missing ART dosing was travelling without sufficient ARV supply and simply forgetting (Fig 2). Longer exposure to ART (aPRR: 0.75, 95% CI 0.64-0.88), p<0.001) and being older (aPRR 0.77, 95% CI 0.76-0.88, p<0.001) were associated with lower risk of detectable viremia. Other variables associated but not statistically significant variables included: being either single, divorced, widowed or separated (aPRR 1.32, 95% CI 0.99-1.78, p=0.061); secondary school level completion (aPRR 1.17, 95% CI 0.85-1.61, p=0.326); travel time from the cART clinic of 1 hour or more (aPRR 0.88, 95% CI 0.66-1.91, p=0.417) and being aware of antiretroviral (ARV) medication dosing instructions (aPRR 1.08, 0.81-1.43, p=0.612). Despite 11% of women not disclosing their HIV status to their male partners, this variable was not significantly associated with detectable viremia. Additionally, about 10% (n=29) of women with unsuppressed viremia had an HIV-uninfected partner.

**Table 2:** Factors associated with detectable viral load >200 copies/ml.

| Variable | Multivariable<sup>1</sup> |  | Multivariable<sup>2</sup> |  |
| --- | --- | --- | --- | --- |
|  | RR (95% CI) | p-value | aRR (95% CI) | p-value |
| Age (5-year increase) | 0.77 (0.67-0.88 ) | <.001 | 0.77 (0.67-0.88) | <.001 |
| Marital status (ref: married/regular partner) |  |  |  |  |
| Other (single, divorced, widowed, separated) | 1.55 (1.19-2.04) | 0.001 | 1.32 (0.99-1.78) | 0.061 |
| Employment (ref: formal employment) |  |  |  |  |
| Not employed | 0.92 (0.65-1.32) | 0.655 | 0.84 (0.58-1.21) | 0.349 |
| Self employed | 1.09 (0.75-1.58) | 0.646 | 1.11 (0.76-1.63) | 0.577 |
| Education (ref: secondary school not complete) |  |  |  |  |
| Secondary school complete | 1.04 (0.78-1.38) | 0.806 | 1.17 (0.85-1.61) | 0.326 |
| Electricity in the premises (ref: Yes) |  |  |  |  |
| No | 1.62 (1.22-2.14) | <.001 | 1.74 (1.28-2.36) | <.001 |
| Tap water in the premises (ref: Yes) |  |  |  |  |
| No | 1.24 (0.96-1.62) | 0.103 | - | - |
| Travel time to clinic from home (ref: less than 1 hour) |  |  |  |  |
| 1 hour or more | 1.01 (0.77-1.33) | 0.936 | 0.88 (0.66-1.19) | 0.417 |
| Condom use during sex in last three months (ref: Never) |  |  |  |  |
| Always | 1.12 (0.74-1.72) | 0.585 | - | - |
| Sometimes | 1.05 (0.71-1.56) | 0.811 | - | - |
| HIV status disclosure to partner (ref=Yes) |  |  |  |  |
| No | 1.45 (0.94-2.25) | 0.096 | - | - |
| Abnormal vaginal discharge in last three months (ref: No) |  |  |  |  |
| Yes | 2.12 (1.32-3.39) | 0.002 | 1.88 (1.16-3.04) | 0.010 |
| Hospital admission in last three months (ref: No) |  |  |  |  |
| Yes | 2.29 (1.20-4.36) | 0.012 | 2.48 (1.28-4.80) | 0.007 |
| Aware of ARV medication dosing instructions (ref: No) |  |  |  |  |
| Yes | 1.05 (0.79-1.41) | 0.723 | 1.08 (0.81-1.43) | 0.612 |
| Missed ART doses since last visit (ref: None) |  |  |  |  |
| Missed some doses | 2.01 (1.55-2.60) | <.001 | 1.63 (1.24-2.13) | <.001 |
| Time since ART initiation<br>(per 1-year increase) | 0.68 (0.57-0.81) | <.001 | 0.75 (0.64-0.88) | <.001 |
| <sup>1</sup>Each predictor fitted separately while adjusted for the country <sup>2</sup>Multivariable model with many predictors |  |  |  |  |

**Fig 2.**
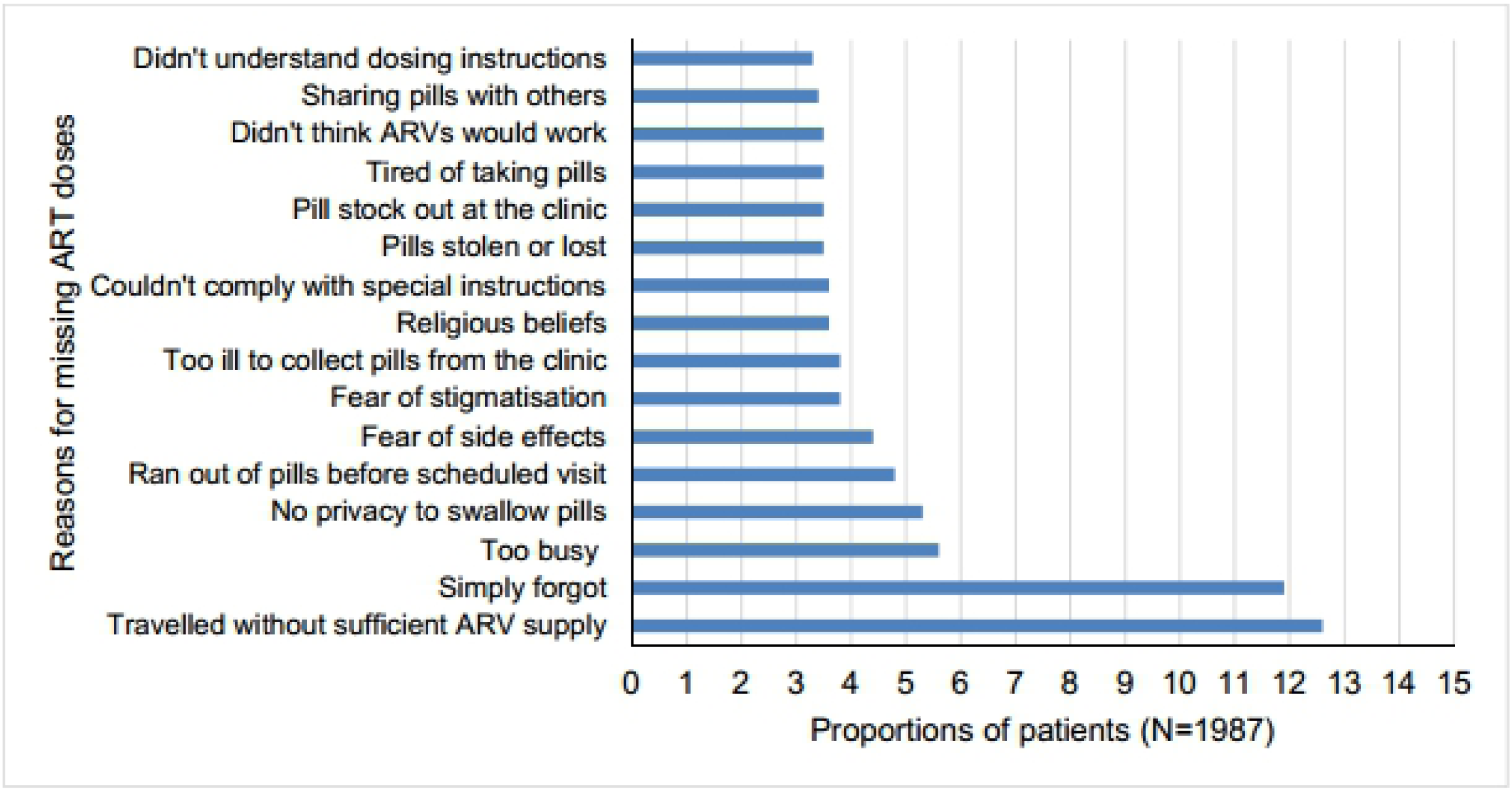
Reasons for missing ART dose.

## Discussion

We found that 11% of the 1934 women initiated on antiretroviral treatment had unsuppressed viremia as defined for the purposes of this study as >200 copies/ml. This is slightly above the UNAIDS 10% target bearing in mind that the 200 copies/ml threshold is lower than the 1000 copies threshold used by UNAIDS. We identify sociodemographic, self-reported non-adherence, and clinical factors that were associated with detectable viremia > 200copies/ml.

The higher proportion of women with detectable viremia than the UNAIDS target suggests that challenges to achieving the 3^rd^ ‘90’ still persist even among women who are in an ideal setting (research) compared to the programs in resource limited settings(3). The association between sociodemographic factors namely the absence of household electricity, a proxy for lower economic status, and detectable viremia above 200 copies/ml, were consistent with other literature that has highlighted that sociodemographic factors are key predictors to poor adherence despite one being on cART(5, 14). The association between younger age and unsuppressed viremia is consistent with current literature (3, 4). Additionally, longer duration of cART use as protective of unsuppressed viremia is consistent with this observation. In contrast to other literature, we found that employment, education and travel time to the clinic were not significantly correlated with viral suppression in the PROMOTE cohort at baseline in the multivariate analyses(14).

Based on self-reported adherence reports using the study questionnaire, we found that self-reported missed ART doses was significantly associated with unsuppressed viremia. Even though there is uncertainty regarding the reliability of self-reported adherence, this measure of adherence still remains as a cheap and easily determined mode of adherence monitoring in resource limited settings using appropriate tools(5). We noted however that a small proportion of women (9.7%) were not aware of the dosing instructions. This factor was not a significant predictor of viral detectability. This finding may suggest that patient education is still lacking. Effective interventions like motivational adherence counseling ensure two-way input and steers away from the traditional methods of adherence counselling(3).

The most common reason for missed doses was travelling with insufficient ARV supply and forgetting, which are in line with other literature(14). Re-emphasis on simple measures for example, setting an alarm (eg mobile phone alarm) and linking dosing with daily activities, should be part and parcel of adherence counseling. Provision of a pillbox is a tool used by participating South African sites. Traveling without ARV supply as a reason could fall in the “forgetting” category, or could be a cover for non-disclosure while visiting family homes. The reason of no privacy, be it at work or in home, also implies non-disclosure and feared stigmatization.. Of note, condom use was not associated with viral detectability above 200 copies/ml.

Relatedly, 11% of all the women had not disclosed their HIV status to their primary partner but this was not predictive of detectable viremia. This PROMOTE result is contrary to results of other studies citing non-disclosure as being associated with poor adherence in different HIV infected populations(14). Contextual issues however remain a major underlying contributor to detectable viremia and patients may not be forthcoming with these reasons for a missed dose. In hyperendemic settings with advanced HIV outreach programmes, one would expect less disclosure issues and united communities taking treatment together, however inherent characteristics prevent effectiveness of the existing adherence promotion programmes.

Other participants provided reasons as being too busy and running out of treatment prior to visit. This is commonly due to life’s every day demands including work commitments. Countries are now working towards improving the access to medicines by means of decentralized dispensing for convenient pill collection. South Africa has implemented a new model where the dispensing services are contracted to private pharmacies(15). Other means of differentiated care strategies to ensure patient convenience include multi-month prescriptions, fast-track refills and community adherence groups who assist with collection and distribution of cART as done in Malawi, Uganda and Zimbabwe(3, 16).

Whereas adherence barriers like fear of side effects, pill burden, perceptions that ART is harmful, feeling sick and depressed have been associated with virologic detectability in various AIDS Clinical Trial Participants in United States, current first-line cART comprises low pill burden and improved safety profile with low toxicity(17). The aim of the health system and its providers is to ensure adherence to first-line cART regimens to prevent the need for second- and third-line cART which are more toxic and have greater pill burden.

More so, some clinical factors like recent hospitalization and abnormal vaginal discharge were significantly associated with viral detectability. Recent hospitalization may suggest clinical failure in this subgroup of non-suppressed women. About 17% of those with detectable viremia had low level viremia of 200 to 999 copies/ml. Other studies have shown that persistent viremia above 200 copies/ml is associated with a higher risk of virologic failure, mortality and morbidity especially among cases of delayed ART switch to 2nd line therapy (18, 19). In addition, 15% of the non-suppressed women were breastfeeding babies born after the PROMISE index child. This alludes to significant adherence challenges which often arise during the postpartum period and beyond(20).

This study contributes much needed data regarding the factors associated with unsuppressed viremia among African pregnant and breastfeeding women receiving ART treatment for life; more so because viral load testing is only a fairly recent intervention in HIV treatment monitoring. These data demonstrate that socioeconomic barriers remain key predictors of viral detectability, as well as recent hospitalization and recent abnormal vaginal discharge.

The strengths in these analyses include that PROMOTE is one of the largest current longitudinal cohort studies of HIV infected women of child bearing age in Africa; and is being conducted in multiple sites in East and Southern Africa, which increases the generalizability of the findings. In addition, there is ongoing Quality Assurance and monitoring as part of the study. Relative limitations to the analyses are that this is a baseline cross sectional analysis; and that data on resistance are not currently available.

### Future plans for the study

Follow up trends in viral load and adherence data over the 5 year follow up in PROMOTE will be presented when available; as will the relation of hair drug levels and drug resistance testing correlates. Point- of- care VL testing coupled with motivation adherence counseling and adherence risk assessment tool development are in the process of being implemented at the sites in Zimbabwe and Uganda respectively.

## Conclusion

This baseline analysis of the PROMOTE study set out to evaluate what clinical and socioeconomic factors were associated with a detectable viremia of >200 copies/mL in African mothers on lifelong cART. Baseline data demonstrate that socioeconomic barriers such as poverty, not being married, young age, and prior history of missing pill doses remain key predictors of viral detectability. This study supports the use of self-reported adherence to cART in the absence of superior adherence measures. The most common reasons given by mothers for missing cART doses emphasize the need for effective motivational adherence counseling, empowering women to improve adherence by using simple reminders and a differentiated care model tailored to mother’s needs. The PROMOTE study insights provide opportunities for possible development and improvement of targeted /cost effective implementation strategies to help support lifetime maternal adherence to both cART and HIV care.

## Disclaimer

The findings and conclusions reported herein are those of the author(s) and do not necessarily reflect the official position of the U.S. government.

## Acknowledgments

We thank the women and children who are participating in the PROMOTE study at each of the research sites. We acknowledge the research teams at each of the following sites: MUJHU, Kampala, Uganda; UNC Project Clinical Research Site, Lilongwe, Malawi; Johns Hopkins-College of Medicine Research Project, Blantyre, Malawi; University of Zimbabwe College of Health Sciences Clinical Trials Research Centre (UZCHS-CTRC) Zimbabwe; Perinatal HIV Research Unit (PHRU), Soweto, South Africa; Centre for the AIDS Programme of Research in South Africa (CAPRISA), uMlazi Clinical Research Site, Durban, South Africa.

## Supporting information

**S1 Manuscript data**

**S2 Variable name and label**

